# Abundance and diversity of the fecal resistome in slaughter pigs and broilers in nine European countries

**DOI:** 10.1101/194647

**Authors:** Patrick Munk, Berith Elkær Knudsen, Oksana Lukjacenko, Ana Sofia Ribeiro Duarte, Roosmarjin E. C. Luiken, Liese Van Gompel, Lidwien A. M. Smit, Heike Schmitt, Alejandro Dorado Garcia, Rasmus Borup Hansen, Thomas Nordahl Petersen, Alex Bossers, Etienne Ruppé, Ole Lund, Hald Tine, Sünje Johanna Pamp, Håkan Vigre, Dick Heederik, Jaap A. Wagenaar, Dik Mevius, Frank M. Aarestrup

## Abstract

**EFFORT group:** Haitske Graveland, Alieda van Essen, Bruno Gonzalez-Zorn, Gabriel Moyano, Pascal Sanders, Claire Chauvin, Julie David, Antonio Battisti, Andrea Caprioli, Jeroen Dewulf, Thomas Blaha, Katharina Wadepohl, Maximiliane Brandt, Dariusz Wasyl, Magdalena Skarzyñska, Magdalena Zajac, Hristo Daskalov, Helmut W Saatkamp, Katharina D.C. Stärk.

**Abstract:** Antimicrobial resistance (AMR) in bacteria and associated human morbidity and mortality is increasing. Use of antimicrobials in livestock selects for AMR that can subsequently be transferred to humans. This flow of AMR between reservoirs demands surveillance in livestock as well as in humans. As part of the EFFORT project (www.effort-against-amr.eu), we have quantified and characterized the acquired resistance gene pools (resistomes) of 181 pig and 178 poultry farms from nine European countries, generating more than 5,000 gigabases of DNA sequence, using shotgun metagenomics. We quantified acquired AMR using the ResFinder database and a database constructed for this study, consisting of AMR genes identified through screening environmental DNA. The pig and poultry resistomes were very different in abundance and composition. There was a significant country effect on the resistomes, more so in pigs than poultry. We found higher AMR loads in pigs, while poultry resistomes were more diverse. We detected several recently described, critical AMR genes, including *mcr-1* and *optrA*, the abundance of which differed both between host species and countries. We found that the total acquired AMR level, was associated with the overall country-specific antimicrobial usage in livestock and that countries with comparable usage patterns had similar resistomes. Novel, functionally-determined AMR genes were, however, not associated with total drug use.

## Introduction

Antimicrobial resistance (AMR) is considered one of the largest threats to human health.^1^ In addition to the use of antimicrobial agents for humans, livestock is considered an important source of AMR, potentially compromising human health.^2^ Besides AMR in zoonotic pathogens, AMR in commensal bacteria is worrisome because of its ability to spread horizontally to pathogens.

Multiple studies have shown that use of antimicrobials in livestock will lead to increased occurrence of AMR and that reduction of usage will eventually lead to reduced resistance.^3^–^8^ A number of national surveillance programs have been implemented to monitor the occurrence of AMR in different reservoirs and follow trends over time.^1^,^9^–^11^ There are major differences in antimicrobial consumption patterns between different countries globally and also within Europe.^12^ Major differences in the occurrence of AMR have also been observed among indicator organisms (e.g. *E. coli*) isolated from different European countries.^3^,^13^ Current monitoring efforts are mainly based on culturing indicator bacteria followed by phenotypic AMR determination.^13^,^14^ This procedure only targets a limited number of species present in the gut microbiota and thus likely represents only a fraction of its resistome (the collective pool of AMR genes). Such metagenomic approaches have been used in a number of recent studies and it has been shown that metagenomic read mapping describes AMR abundance in bacterial communities more accurately than commonly used technologies on selected indicator organisms.^15^–^17^ A recent study focused on sampling a diverse group of individual pigs from eleven farms in three countries showed that genetics, age, diet and country all likely influence the pig microbiota, but little information is available for poultry.^16^

As part of the EU-funded EFFORT project (www.effort-against-amr.eu), we sampled over 9000 animals in 181 pig and 178 poultry herds in nine European countries, generating herd-level composite samples as previously described.^17^ This gives us a unique insight into the diversity and structure of the acquired pig and broiler resistomes across Europe. We sampled animals as close as possible to slaughter to elucidate the potential consumer exposure to AMR associated with meat production in Europe. Association between AMR gene abundance and country level antimicrobial usage was analyzed. We hereby provide an overview of AMR in the two most intensively raised European livestock species. To our knowledge, this study represents the single largest metagenomic AMR monitoring effort of livestock: both in terms of countries (9), herds included (359), individual animals sampled (over 9,000) and sequencing effort (5,000+ gigabases).

## Methods

### Farm selection and sampling

The sampling protocol for pig and broiler farms that has been agreed on by the EFFORT consortium is described below. Selection of farms and sampling procedure followed these guidelines to the extent possible, but some deviations from the protocol were occasionally necessary. A detailed description of the sampling conducted in the individual countries is provided in supplementary material.

### Selection of pig and poultry farms

In each participating country, 20 conventional integrated pig farrow-to-finisher non-mixed farms were selected. The farms needed to have a minimum of 150 sows and 600 fatteners and employ batch production to ensure that the majority of the animals of the sampled group originated from the same birth cohort. All-in all-out production at compartment level was preferred and all fatteners sampled were required to have been on the same site during their entire life. Selected farms should have no contact through livestock trade, and have a random regional distribution.

In each country, 20 conventional broiler farms (no breeders) were selected. The farms should have all-in all-out production, with a thinning procedure from day 30 onwards allowed. All selected farms should have no intended slaughter age higher than 50 days, no slow growing breeds (intended growth rate less than 55 gram/day) and no stocking density lower than 10 birds/m^2^. Only one flock/house per holding should be sampled and the flock should be between 20,000 and 40,000 birds. If possible, selected farms should have a random regional distribution.

### Procedure for sampling

We sampled pig farms between May 2014 and December 2015, and tried to minimize seasonal influences. The sampled fatteners should be as close to slaughter as possible (i.e. within the last week). A total of 25 fresh, still warm and undisturbed fecal droppings were sampled from pen floors (a minimum of 10 g of feces per sample) randomly divided over all eligible compartments/stables of fatteners close to slaughter.

Broilers were sampled between May 2014 and June 2016, and we tried to minimize seasonal influences. On each farm, 25 undisturbed, fresh main bowel droppings were collected from the floor of the house (a minimum of 3 g feces per sample). The flocks should be sampled as close to slaughter as possible (last week before the final depopulation).

All samples were collected aseptically in plastic containers and were stored at 4°C and transported to the laboratory within 24 hours after sampling.

### Pooling and handling of samples

Upon arrival in the laboratory, individual fecal samples were homogenized by stirring thoroughly with a sterile tongue depressor/ spoon for a few minutes. From each pig sample, two 2 ml cryotubes were filled and frozen immediately at -80°C (alternatively at -20°C for maximum 4 days, before transferring to -80°C). For broiler samples, at least 0.5 g feces was added to two cryotubes. Sample pooling was either done immediately or the frozen tubes were shipped to the Technical University of Denmark (DTU) on dry ice for pooling. Individual samples from the same herd were defrosted and placed on ice briefly before weighing. Following weighing, they were pooled with 0.5 g of feces from each sample and stirred for a few min with a sterile device (e.g. disposable wooden tong depressor). All samples were only thawed once shortly before DNA extraction.

After removing two mis-labeled samples, we ended up with composite fecal samples from 178 broiler flocks and 181 pig herds.

### Sampling to estimate the effect of random sampling

To study the potential effect of sampling randomness and the reproducibility of our sampling protocol, two of the pig herds were chosen for triplicate sampling. These two herds were sampled three times on the same day (25 samples x 3 sampling rounds), resulting in six pooled samples (2 herds x 3 sampling rounds), from which the within-farm variation was assessed. A table with all samples and their metadata is included as Supplementary Table 1.

### DNA extraction and sequencing

From each of the pooled, herd-level fecal samples, DNA was extracted using a modified QIAamp Fast DNA stool mini kit protocol (Qiagen, cat. no. 51604), as previously described.^18^ One major modification is the addition of a bead beating step in the beginning of DNA extraction. The protocol can be found at https://figshare.com/articles/SOP_-_DNA_Isolation_QIAamp_Fast_DNA_Stool_Modified/3475406. DNA-purification of all pooled samples was processed centrally at Technical University of Denmark (DTU), and the DNA was stored in duplicates at -20°C until further use.

DNA was shipped on dry ice for library preparation and sequencing at the Oklahoma Medical Research Foundation (OMRF). There, DNA from all samples was mechanically sheared to a targeted fragment size of 300bp using ultrasonication (Covaris E220evolution). For pooled pig samples, library preparation was performed with the NEXTflex PCR-free library preparation kit (Bioo Scientific). For poultry samples, due to a lower DNA availability, the minimal amplification-based KAPA Hyper kit (Kapa Biosystems) was used. For all samples, the Bioo NEXTflex-96 adapter set (Bioo Scientific) was used. In batches of roughly sixty samples, the libraries were multiplexed and sequenced on the HiSeq3000 platform (Illumina), using 2x150bp paired-end sequencing per flow cell. A total of 17 Belgian, Danish and Dutch pig fecal samples were sequenced on the HiSeq2500 platform (Illumina), using 2x100bp paired-end sequencing, before converting to HiSeq3000 for the remaining samples (See Supplementary Table 1).

### Bioinformatics processing

The FASTQ files with sequencing data for each sample were analyzed following the principles from the previously described MGmapper tool.^15^ The reads were first cleaned by cleaning out adaptors using BBduk (BBMap v39.92 - Bushnell B. -https://sourceforge.net/projects/bbmap/), and by removing reads that aligned to the internal sequencing control phi-X174 as determined by the BWA-MEM algorithm.^19^ Trimmed read pairs were aligned using the BWA-MEM algorithm Prokaryotic RefSeq genomes from NCBI GenBank (downloaded on Nov. 18, 2016) with the “reference” or “representative” genome tags. The BWA-MEM algorithm (v0.7.15) was slightly modified such that the insert size was estimated for the entire sample unaffected by the number of CPU cores used. This estimate was then used for re-analyzing the cleaned read pairs using BWA-MEM, ensuring that the thresholds for a read pair to map as a ‘proper pair’ were the same for all read pairs, and avoiding bad insert size estimates when few read pairs aligned.

The read pairs were aligned to the bacterial NCBI genomes again, and to the AMR genes present in the ResFinder database (accessed Nov. 17, 2016).^20^ ResFinder is a manually curated database of horizontally acquired AMR genes and thus does not include intrinsic AMR genes and mutated housekeeping genes providing AMR by changing the drug target. Technical duplicate read pairs were then removed using ‘MarkDuplicates’ from the Picard command line tools (v2.8.3; http://broadinstitute.github.io/picard/).

For each ResFinder reference sequence, we counted the number of read pairs that properly aligned with at least 50 bp aligning from both the forward and reverse reads. Each read pair matching a ResFinder reference was assigned to the first highest-scoring reference, as done in MGmapper. The same was done for NCBI microbial genomes in order to quantify the bacteriome and get a measure of the microbial proportion within each sample. This total was used to normalize the ResFinder counts, by computing an FPKM-value (fragments per kilobase reference per million bacterial fragments) for each ResFinder reference sequence. The FPKM values were computed by dividing the mapping count on each reference with its gene length and the total number of bacterial read pairs for the samples and multiplying by 10^9^.^21^ Raw mapping count data and their associated FPKM values can be found in Supplementary Table 2 and 3.

Because ResFinder contains many representatives of certain gene families, a high degree of homology exists, with long stretches of the references being identical. This causes unspecific mapping between high-identity sequences. To eliminate this random noise, we chose to aggregate read counts and relative abundances post-mapping at higher levels based on sequence identity. We clustered all the ResFinder genes using CD-HIT-EST (v4.6.6) at a 90% identity level and otherwise default settings.^22^ The resulting gene clusters were manually inspected and named to reflect their contents while avoiding conflicts with other clusters (Supplementary Table 4). Abundances were aggregated according to these clusters of high-identity genes and the resistance-class-level as annotated in ResFinder. These two levels, “gene” and “class”, were used for all downstream analysis.

### Functional resistance database

Previous studies have identified a wide array of novel AMR genes in various reservoirs using functional metagenomics, referred to as functional AMR genes.^23^–^26^ By cloning random DNA fragments from complex microbiomes into an expression vector expressed in a host (typically *E. coli*) and selecting for growth in the presence of certain antibiotics, they have been found to provide AMR to many antibiotics.^23^–^26^ We constructed a functional resistance database (FRD) from 3,416 AMR gene variants identified in four major studies, using 23 different antimicrobials for selection.^23^–^26^

Briefly, in each of these studies, DNA was extracted from environmental and human fecal samples, fragmented and cloned into a plasmid vector and screened for AMR functionality in *E. coli* cultured with one of multiple antimicrobials. AMR-granting plasmid inserts were then sequenced and the responsible open reading frame was identified. The protocol for the database construction can be found at https://cge.cbs.dtu.dk/services/ResFinderFG/. Genes were quantified using MGmapper as was done for ResFinder. Genes with more than 90% identity to ResFinder genes were removed post mapping to obtain the new AMR genes without overlap with ResFinder. The resulting data was aggregated to 90% gene clusters, using CD-HIT-EST, as was done for ResFinder.^22^ The most frequent gene clusters remaining were derived from genes selected using: trimethoprim, chloramphenicol, co-trimoxazole, cycloserine, amoxicillin, gentamicin, penicillin and tetracycline.

### Principal coordinate analysis and resistome clustering

For principal coordinate analysis (PCoA), the gene-cluster level FPKM matrix was Hellinger-transformed and the Bray-Curtis (BC) dissimilarities between all samples were calculated using the R package *vegan*.^27^ PCoA was carried out for both pigs and poultry; combined and separately, using the *vegan* function ‘betadisper’. The same analysis was used to test whether host animal and country were significant predictors of within-group beta diversity dispersion. The effects of country on sample dissimilarities was determined using ‘permutational multivariate analysis of variance using distance matrices’ (‘adonis2’ function in *vegan* package), separately for pig and poultry.

### Antimicrobial use in livestock

Data for national livestock antimicrobial usage (AMU) was obtained from the European Medicines Agency’s 2014 European Surveillance of Veterinary Antimicrobial Consumption (ESVAC) report and was stratified by major drug family.^28^ The mass of active compound sold for use in animals in 2014 was divided by the Population Correction Unit (PCU) in 10^6^ kg - approximating the biomass. The PCU is a unit that allows inter-species integration by adjusting for import/export and differences in average weight between species when they are most likely to receive antimicrobial treatment. The estimate was multiplied by 1000 to obtain drug mg/kg PCU livestock. The country-specific veterinary drug use can be found in Supplementary Table 5.

### Procrustes analyses

In order to test the association between country-specific AMU patterns and the resistomes, we performed Procrustes analysis using the *vegan* R package as follows. A PCoA was generated from Euclidean distances between the samples in the PCU-corrected AMU (Supplementary Table 5, Supplementary Figure 1). The AMU PCoA was tested against the previously mentioned Hellinger-transformed, resistome BC dissimilarity PCoA using the ‘protest’ function with the default 999 permutations, again separately for pigs and poultry.

The gene-cluster FPKM ResFinder matrix and the genus-level FPKM taxonomy matrix were Hellinger transformed and BC dissimilarities were calculated. They were ordinated using non-metric multidimensional scaling (NMDS) with the ‘metaMDS’ *vegan* function (999 permutations) for pig and poultry samples separately. The symmetric Procrustes correlation coefficients between the bacteriome and resistome ordinations, p-values and plots were obtained using the ‘protest’ and ‘procrustes’ functions in *vegan*.^29^

### Alpha diversity

For all samples, we computed the within-herd resistome diversity using Simpson diversity index (1-D), Chao1 richness estimate and Pielou’s evenness.^30^ The raw read count matrix was rarified to 10,000 hits for all samples for alpha diversity estimation, leading to the exclusion of 10 samples.

### Visualization

Heatmaps were produced using the *pheatmap* R package. For heatmaps showing individual gene abundances, the BC dissimilarities between samples were used. For all other dendrograms, the Pearson product-moment correlation coefficients (PPMCC) were used. Complete-linkage clustering was used for all dendrogram clustering. For sample similarities, BC dissimilarity was converted to a similarity percentage, i.e., 100*(1-BC).

The circular BC sample dendrogram was exported in Newick format using the *ape* package and further annotated with the Interactive Tree of Life tool.^31^,^32^ Bar-, box- and scatter plots were produced using the *ggplot2* R library.^33^ The R library *RcolorBrewer* was used to generate the color palettes used. The library is based on work by Cynthia A. Brewer (www.ColorBrewer.org).

### Statistical analyses

All statistics was done in Microsoft R Open (MRO) 3.3.2, using the libraries and procedures detailed below. Exact package versions can be found here: https://mran.revolutionanalytics.com/snapshot/2016-11-01/bin/windows/contrib/3.3/. For statistical tests, only the first sampling from triple-sampled herds was included (see Supplementary Table 1). Unless otherwise mentioned, all statistical analyses were performed on pigs and poultry separately.

### Effect of AMU on total AMR

For testing the effect of total AMU on total metagenomic AMR abundance (sum of all genes), we used the *lme4* 1.1-12 package to make linear mixed effects regression models with total livestock drug usage as the independent variable, total AMR abundance (FPKM) as the dependent variable and country as a mixed effect intercept, adjusting for the fact that AMR abundance observed in farms from the same country is correlated due to factors that are not included in this study.^34^ The effect of AMU was modelled on a logarithmic scale, which resulted in lower Akaike’s information criteria compared with modelling AMU on a linear scale. Country-and sample-level residuals were plotted and inspected for normality and homoscedasticity. Pig sample residuals and country residuals were normal and so were poultry country residuals. Poultry sample residuals had a longer right tail. Square-root transforming the poultry AMR data, gave more normal residuals and a similar conclusion (p<0.05). The effect and significance of drug usage was assessed using likelihood-ratio tests, comparing the random effect models with and without the AMU effect.

### Differential abundance analysis

To identify differentially abundant AMR genes per country, we analyzed the read pair mapping count matrix using the *DESeq2* package as previously recommended for metagenomic read count data.^35^,^36^ This was done on the raw read pair matrix, following recommendations that rarefying is not warranted in metagenomic studies.^36^ The read-pair count matrices for pigs and poultry were analyzed separately. The number of mapped bacterial pairs was divided by the minimum number of mapped bacterial pairs and was used as the size factor. For each gene, we used a two-sided Wald test to determine whether the fold change between countries differed from 0 and extracted all the country-versus-country results. P-values were adjusted for the false discovery rate (FDR) using the Benjamini-Hochberg approach and we used a significance threshold of alpha: 0.05.^37^

### Core resistome

The core resistomes determined here, were the set of AMR gene clusters with mapping read pairs in at least 95% of the samples. The core resistomes were determined separately for the pig and poultry reservoirs.

## Results

In total, DNA from 365 pooled samples was extracted and shotgun sequenced, resulting in more than 36 billion sequences (18 billion paired-end (PE) reads), comprising more than 5 terabases of DNA. This yielded an average of 50 million (SD: 18*10^6) PE reads per pooled sample. This was similar for pig and poultry samples, though the former varied more than the latter.

### Acquired resistome characterization

The total AMR gene level varied significantly across samples, both depending on host animal and country of origin. In general, pigs had a higher AMR level than poultry (Figure 1a). The highest AMR levels were found in Italian pigs from where the top four resistance-scoring samples originated, all in excess of 10,000 FPKM AMR. At the lower end of the spectrum were Danish poultry samples that occupied the eleven samples with least AMR, all below 500 FPKM.

**Figure 1.**
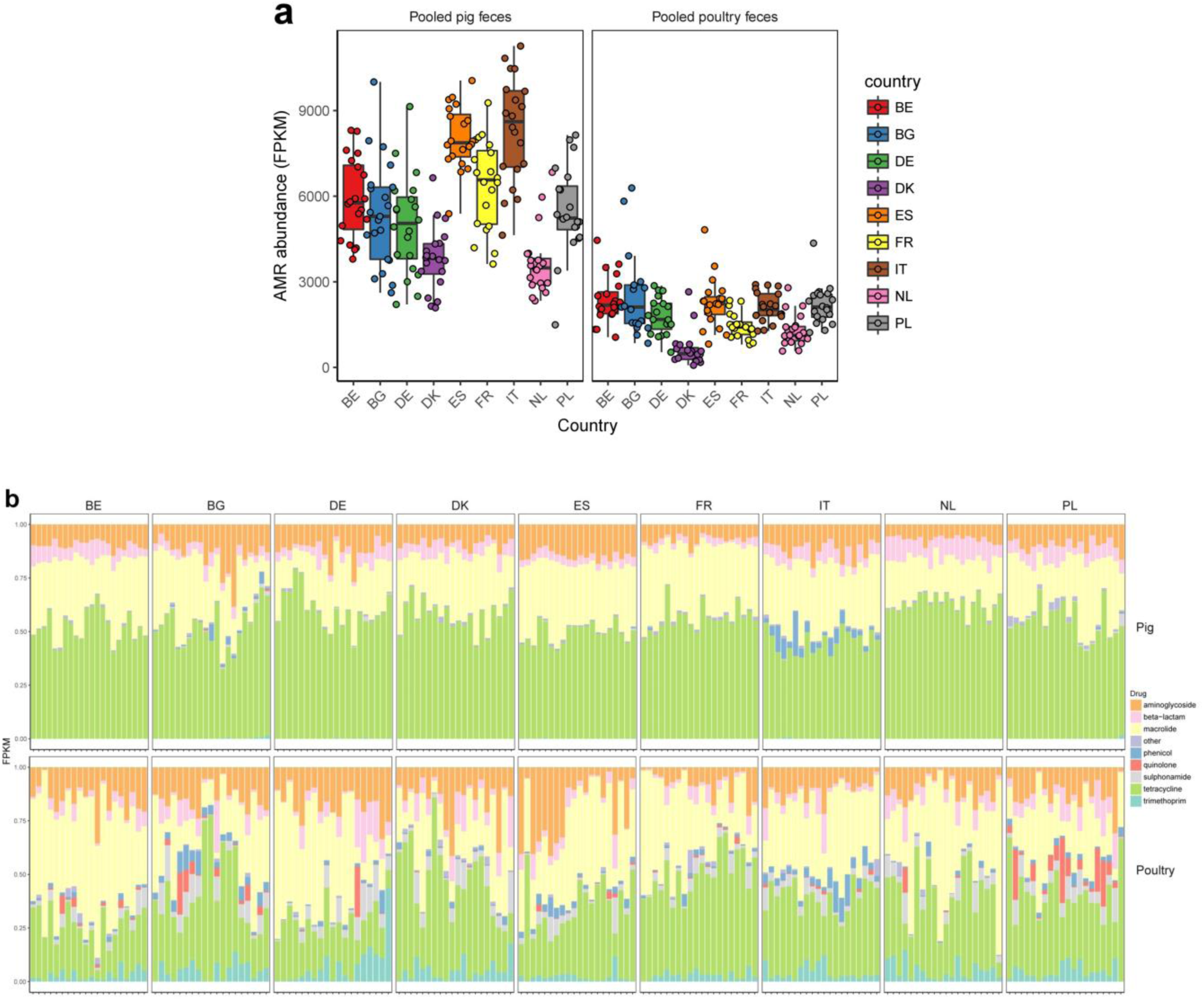
Overview of AMR abundance and composition. From the read mapping results to the ResFinder database, AMR abundance was calculated for each reference gene in each sample. (a) Boxplots showing the total AMR level per sample, stratified by host species and country. Each herd is also represented by a dot with sideways jitter to minimize overplotting. Horizontal box lines represent Q1, median and Q3. Whiskers denote range of points within Q1-1.5*IQR and Q3+1.5*IQR. (b) Stacked barchart of AMR abundance per type (colors) per sample (x-axis), proportional to total AMR within each sample.

We summed the relative abundance of AMR to the corresponding drug class level for each sample to look for major trends across species and countries (Figure 1b). When considering the proportion of the total resistome that each AMR type takes up, the pig samples were relatively homogenous: tetracycline AMR was by far the most common, followed by macrolide AMR. Beta-lactam and aminoglycoside AMR genes followed with other kinds of AMR being rare. Italian pigs had a notably larger proportion of phenicol AMR compared with other countries and it seemed to be consistent across all Italian farms. A subset of Bulgarian pig farms had a similar proportion of phenicol AMR.

Among the poultry farms, there was less consistency: both within and between countries, the relative proportions of AMR per drug class varied more. Tetracycline, macrolide, beta-lactam and aminoglycoside AMR made up the majority of AMR, as in pig samples, but the two latter classes had very minimal contributions in a subset of herds. Sulfonamide and trimethoprim AMR were more abundant in poultry samples compared with pig samples, across all countries. In many Polish poultry herds, quinolone AMR made up a sizeable fraction of the combined resistome. This was also true for a few non-Polish herds, notably in Bulgaria. For non-proportional graphical representations of the AMR load stratified by sample and drug class, see the supplementary material for an unscaled, stacked bar chart (Supplementary Figure 2) and a heatmap (Supplementary Figure 3). Class level AMR relative abundances can be found in Supplementary Table 6.

To characterize the individual components of the resistome, we summed relative abundance to the gene-cluster level as we had done at drug class level. We found evidence for 407 different gene clusters across all pig and poultry samples (Supplementary Table 2).

We calculated the BC dissimilarities between all samples’ gene-level resistomes and visualized it in a dendrogram (Figure 2a). There was a perfect host separation, with all pig samples clustering separately from all poultry, suggesting pig and poultry resistomes are very different. Within the pig cluster, the country separation was more pronounced than for poultry. An exception was Danish poultry, where 18/20 samples clustered.

To assess the reproducibility of our protocol, from sampling through sequencing, we evaluated the similarities between resistomes of two triple-sampled swine herds. Dutch triple-sampled herd pools had the highest similarities of all samples, ranging from 93.6% - 93.7% BC similarity. The Belgian triple-sampled herd pools had values ranging from 91.5% - 93.3% similarity. No replicated sample pool had higher similarity to other herds than to its own replicates and the two sets of three samples can thus be seen clustering as expected (Figure 2a). A sample similarity heatmap is found in Supplementary Figure 4.

**Figure 2.**
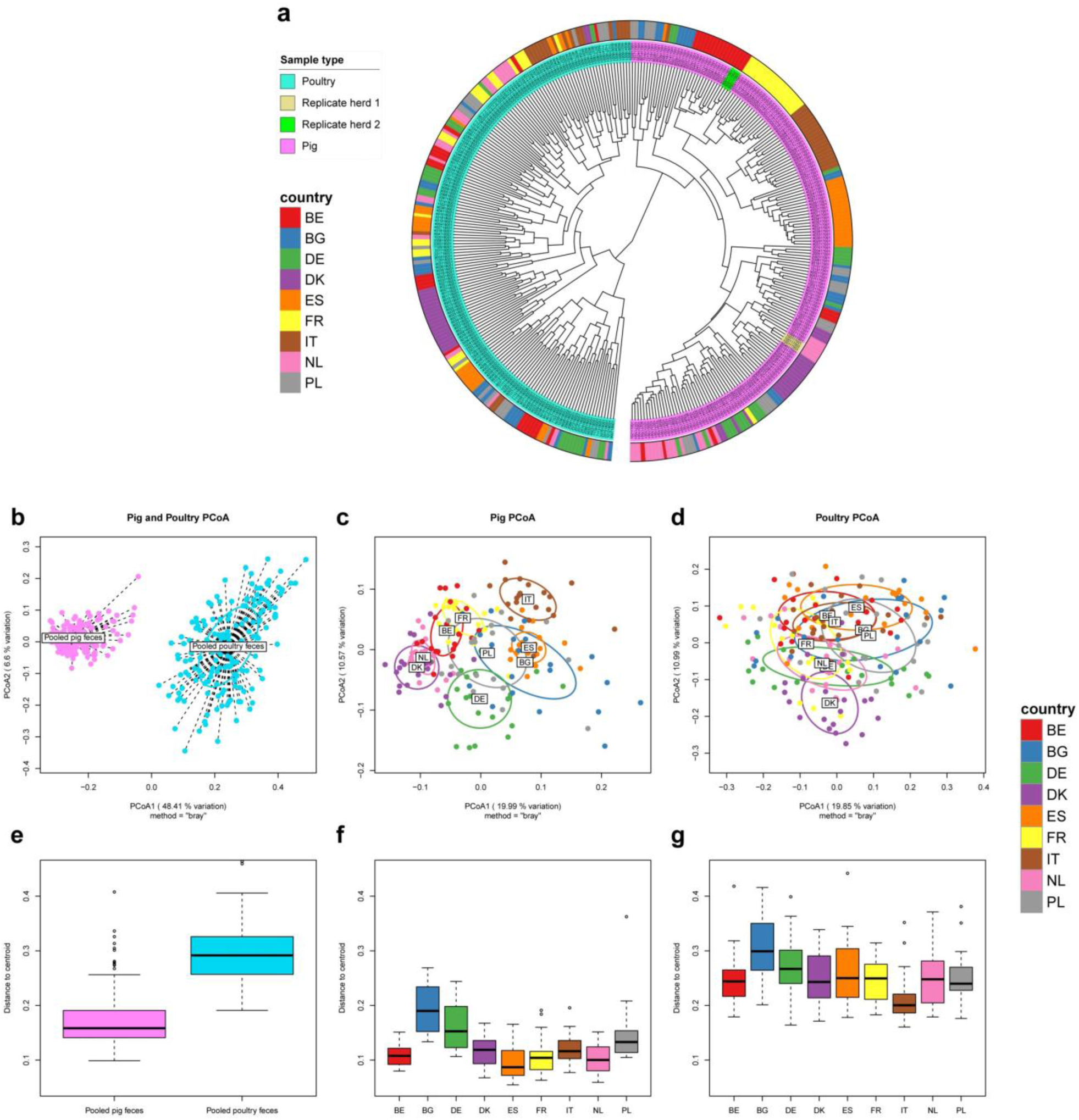
Resistome clustering is influenced by both host animal and country. (a) Dendrogram showing complete linkage clustering of BC dissimilarities between all pig and poultry resistomes. Triple-sampled pig herds are highlighted in separate colors. (b-d) PCoA plots for pig and poultry samples combined (b), pig samples (c) and poultry samples (d), respectively. Ellipses denote standard deviation for distance of each member to its group centroid (labeled). (e-g) Boxplots of distances for each group’s samples to its centroid for pig and poultry samples combined (e), pig samples (f) and poultry samples (g), respectively. Horizontal box lines represent Q1, median and Q3. Whiskers denote range of points within Q1-1.5*IQR and Q3+1.5*IQR.

We ordinated the gene-level resistomes for all samples (Figure 2b) and pig and poultry samples separately (Figure 2c-d). As with hierarchical clustering, there was a clear separation of pig and poultry samples, along the first principal coordinate, which explained 48% of the variation across all resistomes.

When analyzing the two reservoirs separately, we observed clustering according to country of origin in pigs (Figure 2c), while clustering was more diffuse for poultry (Figure 2d). We tested for the country effect and found it to be significant in both pigs (adonis2 p<0.001) and poultry (adonis2 p<0.001). In poultry however, the country effect only explained roughly a quarter of the variation, while country explained roughly half of the variation in pigs (data not shown). In the pig resistome ordination, the Danish and Dutch samples clustered closely together. The same could be seen for French and Belgian resistomes and to a lesser degree, Italian and Spanish samples. Bulgaria, Germany and Poland showed larger dispersions than the other countries. Poultry samples had a higher dispersion than pig samples (Figure 2e, beta-dispersion p<0.001). Beta-dispersion levels varied significantly between countries in both pigs (Figure 2f, beta-dispersion p<0.001) and poultry (Figure 2g, beta-dispersion p<0.001).

We visualized the AMR gene abundances in a heatmap to look at the overall structure and composition of the resistomes and the co-occurrence of AMR genes (Supplementary Figure 5). Some AMR genes were more abundant in one species, while others, including *tet(W)* and *erm(B)* were ubiquitous in all samples, for both species. Among the pig samples, the Italian samples stood out: several chloramphenicol AMR genes, including *cat(pC194)*, *catP*, and *cat_2*, were much more abundant in ltaly, compared to the other countries, consistent with our inspection of AMR at class level (Figure 1). Several AMR genes known to be co-located indeed co-occurred across samples. The genes in the vancomycin AMR *VanA* cassette were co-located in a number of poultry samples. This was also true for the *VanB* cassette members, clustering together, but separately from *VanA*, showing ability to distinguish variants of homologous genes. As earlier indicated, the poultry samples showed less country-based clustering than pigs. An exception were the Danish poultry samples. These had noticeably lower abundance of many AMR genes that were widespread in other countries.

### Core resistome

To determine whether specific genes were unique to each of the host animals, we examined the set of AMR genes that was consistently observed within each animal species (evidence for it in 95% of samples). We identified 33 core AMR genes in pigs and 49 core AMR genes in poultry, with 24 being shared between the two hosts (Supplementary Figure 6). Hence, only nine AMR genes were pig-core genes without also being poultry-core genes. These included the genes making up the Van-G vancomycin cassette, *tet(C)*, *bla*_ACI_ and *cfxA*. Twenty-five AMR genes were poultry-core genes without also being pig-core genes and include the Enterobacteriaceae-associated *strAB*, *sul2*, *blaTEM* and *tet(A)* genes.

### Differential abundance analysis

In order to test which specific genes differed in abundance between countries, we carried out a differential abundance analysis for ResFinder gene read pair counts. Heavy overrepresentation of low unadjusted p-values indicated a large effect of country in both in the pig and poultry datasets (Supplementary Figure 7). Of special interest was the newly characterized *Enterococcus*-associated linezolid-resistance gene *optrA*, that had a significantly higher abundance in Bulgarian poultry farms, compared with poultry farms in all other countries (FDR < 0.05) (Figure 3b). A single Spanish farm did, however, have even higher *optrA* abundance than any other farm. Among the pig herds, the *optrA* gene was more abundant in Bulgarian and Italian herds than anywhere else (except for two farms in Spain) (FDR<0.05).

The newly identified colistin-resistance gene *mcr-1* was significantly more abundant in Bulgarian and Italian poultry farms, compared with most other countries (FDR<0.05). France, Poland and Spain had intermediate levels, while Denmark, the Netherlands and Germany had the lowest levels (Figure 3b).

As previously noted from visual inspection of heatmaps, multiple chloramphenicol AMR genes including *cat(pC194)* were much more abundant in Italian pigs than other pigs. The ESBL *bla*_CTX-M_ gene cluster 1 also showed country dependency, being significantly more abundant in poultry samples from Spain, Poland, Italy, France and Belgium than Germany (FDR<0.05). Differential abundance analysis results can be found in Supplementary Table 7 and 8 for pig and poultry respectively.

**Figure 3.**
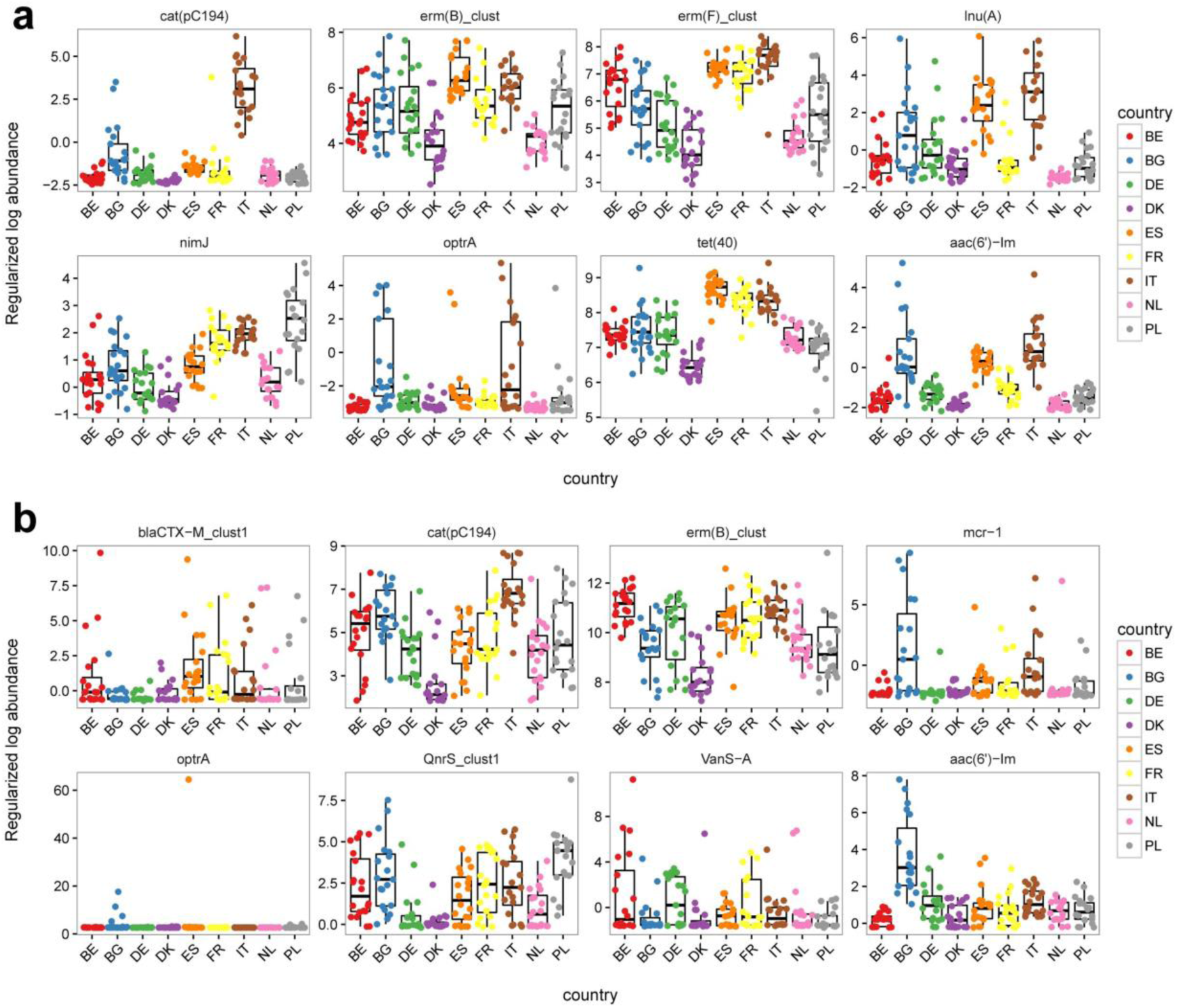
AMR genes differ in abundance between countries. A handpicked subset of genes that differed significantly in abundance between at least two countries’ pig farms (a) or poultry farms (b). The regularized log abundance (rlog) is shown on the y-axis in boxplots and points. Points were sideways jittered to reduce overplotting. Horizontal box lines represent Q1, median and Q3. Whiskers denote range of points within Q1-1.5*IQR and Q3+1.5*IQR.

### Alpha diversity and richness

We calculated several alpha diversity indexes for each farm resistome (Figure 4) The range of AMR diversity was generally much larger for poultry samples, having both lower and higher diversity than pig samples, which had a tighter spread of diversity. The poultry samples had a higher estimated richness than pigs (i.e. a higher number of unique AMR genes per sample). Alpha diversity indexes can be found in Supplementary Table 9.

Interestingly, countries with high AMR richness in pigs also tended to have high AMR richness in poultry. Spain had the highest median richness in both reservoirs, followed by Italy. Poland and Bulgaria together had the third and fourth highest AMR richness in pigs and poultry. Overall, the median estimated Chao1 richness per country correlated significantly between the reservoirs (Spearman’s rho: 0.88, p < 0.01). This was neither true for evenness nor diversity (p>0.05). Rarefaction curves for pig and poultry resistomes can be found in Supplementary Figure 8.

**Figure 4.**
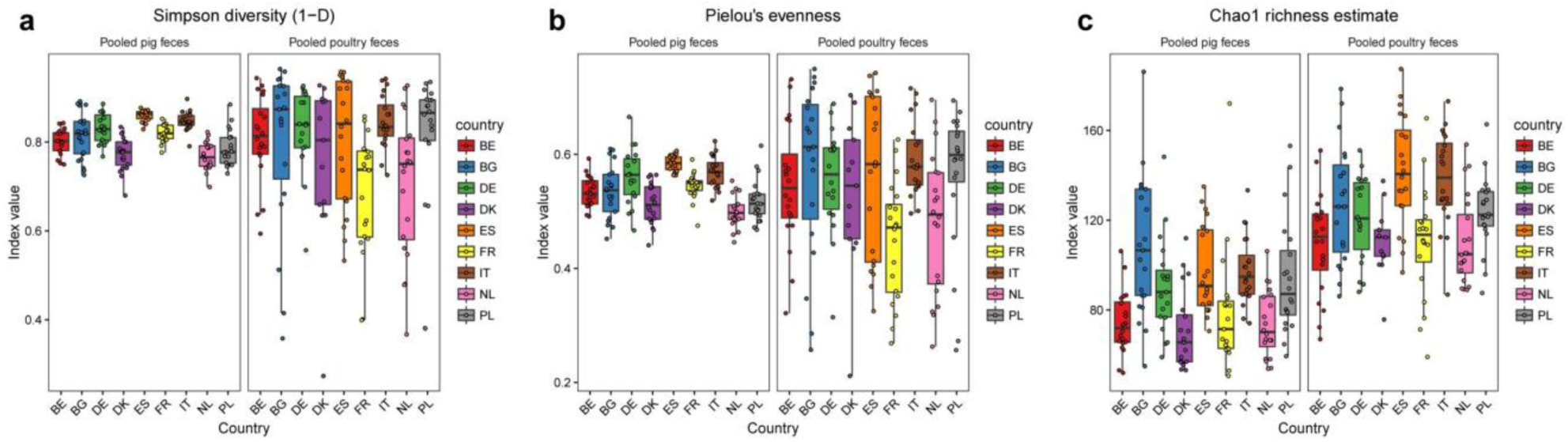
Resistome alpha diversity and richness differs between animal host and countries. From the read count pair matrix, several indexes were calculated. (a) Simpson diversity index, (b) Pielou’s evenness, and (c) the Chao1 richness estimate. Horizontal box lines represent Q1, median and Q3. Whiskers denote range of points within Q1-1.5*IQR and Q3+1.5*IQR.

### Association between bacteriome and resistome

To test the degree to which bacterial genus composition of the microbiota dictates the resistomes, Procrustes analyses were performed. We compared ordinations of the pig microbiome with the pig resistome and the poultry microbiome with the poultry resistome (Figure 5). We found that for both pig and poultry, the bacterial composition correlated significantly with the resistome (p<0.001). Samples with similar taxonomic compositions thus tended to have similar resistome compositions.

**Figure 5.**
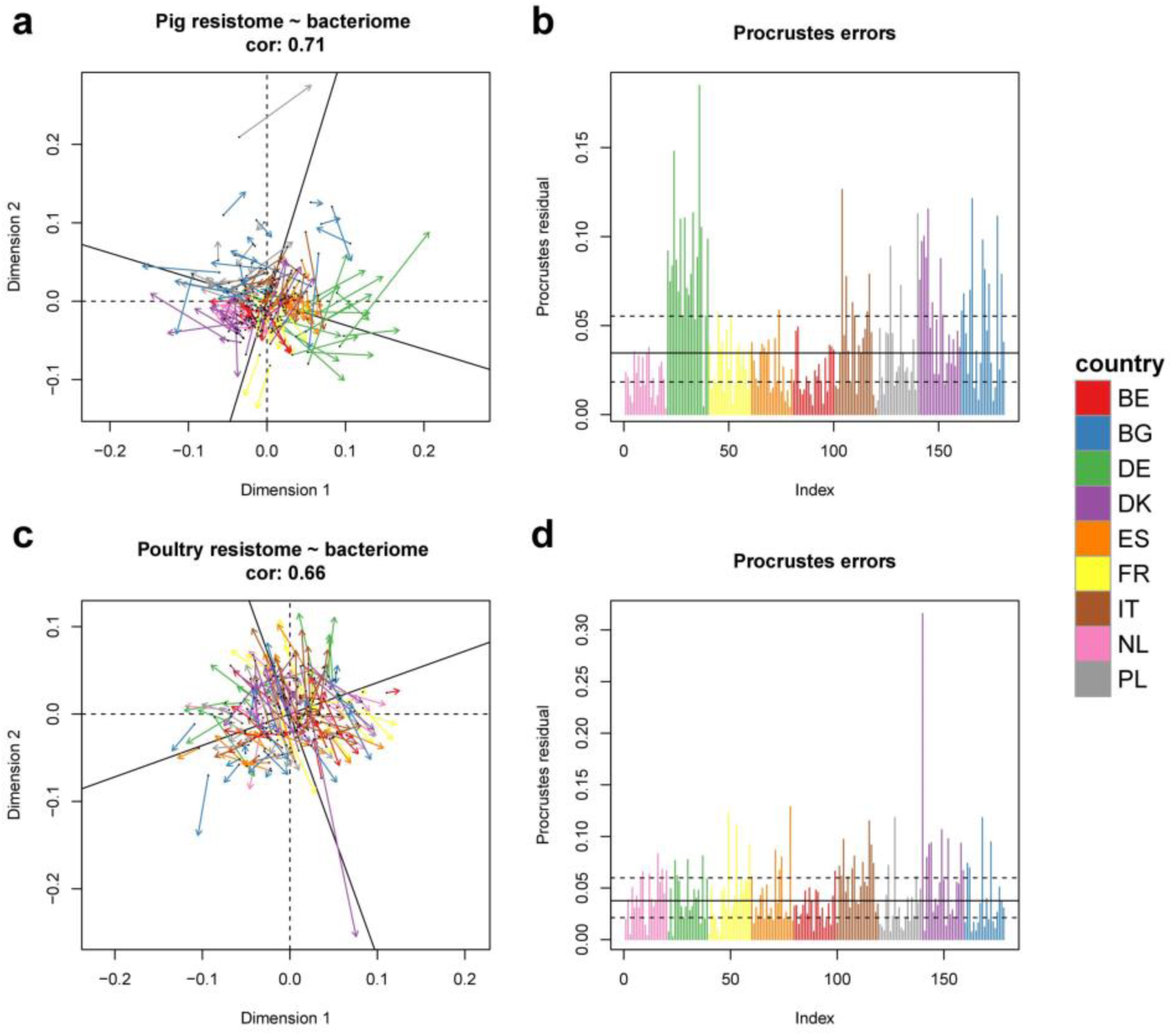
Microbial composition dictates the resistome. Procrustes analysis plots for pig (a/b) and poultry (c/d). Procrustes rotations of bacterial composition onto their corresponding resistome composition (a/c). Bar charts of Procrustes residuals (b/d).

The correspondence between the two datasets was slightly stronger in pigs (Procrustes symmetric correlation: 0.71) than in poultry (Procrustes symmetric correlation: 0.66). Interestingly, in pig samples we saw a country effect on the strength of association between the bacteriome and the resistome. In Dutch and Belgian pig herds, ordinations based on bacterial genera and AMR genes gave similar results (Figure 5b). For samples from Bulgaria, Denmark, Italy and especially Germany however, the Procrustes residuals were larger. This was less evident for poultry, though a single Danish poultry herd had a very unusual resistome, considering its taxonomic composition (Figure 5d). Stress plots for NMDS can be found in Supplementary Figure 9.

### AMR and drug use association

We found that the total country-level veterinary AMU was positively associated with AMR in both pigs and poultry. The AMR abundance increased by 1736-3507 (95% CI, β=2621) FPKM in pigs when the AMU increased by 1 log_e_ unit (36.8% increase in AMU) (Figure 6a) and to a lesser degree in poultry, where the AMR abundance increase by 68 - 1330 FPKM (95% CI, β =700) when the AMU increase 1 log_e_ unit (Figure 6b). For pigs, the variance between farms within countries was 7 times larger than the variance between countries in general, whereas in poultry the variance was 4 times larger within-country than between countries.

**Figure 6.**
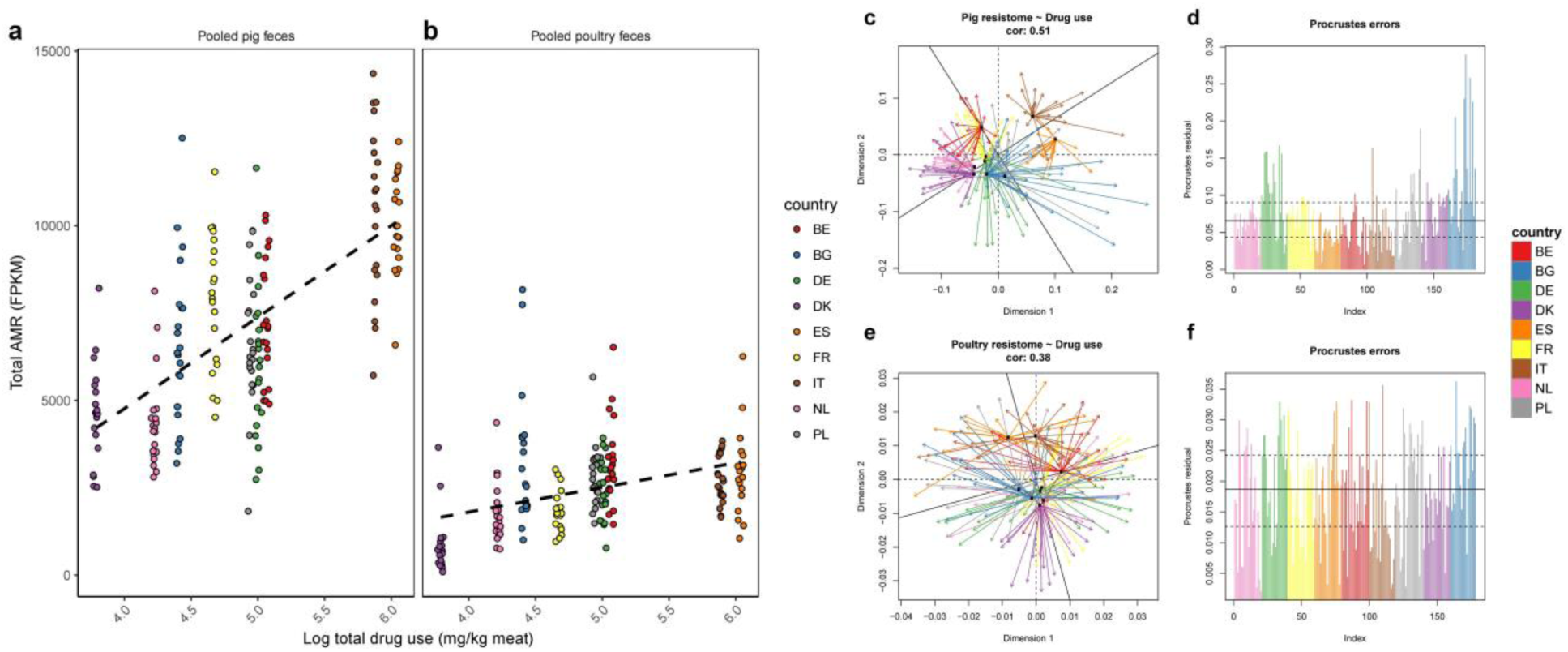
National veterinary AMU affects total metagenomic AMR. (a-b) Scatter plots of country AMU and pooled sample total AMR. A slight sideways jitter was added to the points to minimize overplotting. Trend lines are shown for (a) pigs and (b) poultry. (c-f) Procrustes errors from rotation of resistome principal coordinate analysis (PCoA) onto country AMU PCoA (c-d) and poultry (e-f). (c/e) Procrustes rotations with arrows starting in AMU and ending in corresponding AMR ordination. (d/f) Line plots showing length of each residual arrow in Procrustes rotation.

To test if the AMU pattern was associated with AMR gene profiles, we compared the AMR abundance matrix with AMU matrix, comprised of the 15 recorded AM classes (Supplementary Table 5). We found an association between the veterinary AMU pattern and the pig resistomes (Procrustes correlation: 0.51, p<0.001) (Figure 6c-d). The poultry resistomes were also significantly associated with AMU, albeit with a lower correlation, likely due to their larger beta-diversity and lower degree of country clustering (Procrustes correlation: 0.38, p<0.001) (Figure 6e-f).

### Functional AMR genes

In addition to using ResFinder, we also ran most analyses with the FRD database, to elucidate whether the functionally determined AMR genes behave similarly to the acquired AMR genes in ResFinder. If FRD genes serve similar AMR functionality as the acquired ResFinder genes, we would expect similar results.

Using the FRD database, we found both similar and different patterns, compared with using ResFinder. There was still a perfect separation between pig and poultry samples, but the country separation in pigs was less distinct than when using ResFinder (Supplementary Figure 10). Though less variation could be explained by two axes, the PCoA plot of pig samples now clustered German and Spanish samples, with the remaining countries being more similar. The resistome richness showed similar patterns to ResFinder: Spanish, Italian, Polish and Bulgarian samples had a higher estimated richness in both pig and poultry, compared to the other countries. The Procrustes correlation between the resistome and drug usage was lower (0.40 for pig and 0.25 for poultry). This result was echoed by the lack of association between total AMR and total AMU, for both pig and poultry (p>0.05, Supplementary Figure 11).

## Discussion

Using a metagenomic shotgun sequencing strategy, we were able to detect and quantify more than 400 AMR genes across 181 pig and 178 poultry herds in 9 European countries.

A recent study including Chinese, Danish and French pigs showed the Chinese pig resistomes clustered separately, while Danish and French overlapped.^16^ Here we demonstrate that even among European countries, the livestock resistomes differ in a country-specific manner that might be explained by differential AMU so that countries with similarly high and diverse AMU (Spain, Italy) have similar resistomes, the same way as countries with similarly low AMU (Denmark, Netherlands) also have similar pig resistomes.

We found that the beta diversity dispersion seems to be country dependent, particularly in pigs, with Bulgarian, German and Polish pig herds having more dispersed AMR. While we cannot currently explain this, we consider possible causes as differences in trade and management, among others.

We found the recently discovered plasmid-borne colistin resistance gene *mcr-1* in a number of poultry herds, especially in Bulgaria, Spain and Italy. Spain and Italy had the highest reported veterinary colistin usage among the surveyed countries, whereas Bulgaria has a low reported usage, uncharacteristic for the high *mcr-1* level found here.^13^ This gene was recently discovered in China and identified throughout the world and has been identified in pigs, poultry and human clinical infections alike.^38^

A newly characterized enterococcal linezolid-resistance gene, *optrA* was detected in a subset of pig samples, with Bulgaria, Italy and Spain having the highest abundances. The *optrA* gene provides AMR to both oxazolidinone and amphenicols, including the veterinary-used florfenicol.^13^,^39^ The high abundance of this gene in these countries can likely be explained by the fact that they have the highest veterinary amphenicol usage (together with Croatia, which we did not sample) among the 26 countries surveyed by ESVAC.^12^ This explanation fits well with the fact that Bulgaria, Italy and Spain also had the highest abundances of chloramphenicol AMR genes such as *cat(pC194)* in poultry.

Another AMR gene of special interest, the *bla*_CTX-M_, was also observed in the poultry herds. The higher abundance of *bla*_*CTX-M*_ cluster 1 in Spain, Italy, Poland and Belgium, could possibly be explained by co-selection by fluoroquinolones, which is used more in Spain, Poland, Italy and Belgium than other sampled countries. *qnr* and *bla*_CTX-M_ genes are frequently co-located on large ESBL plasmids. Veterinary cephalosporin usage did not seem to explain the observed levels.

Poland and Spain use far more fluoroquinolones veterinarily than other countries included in this study. We found that plasmid mediated quinolone AMR (*qnr*-genes) was frequently abundant in Polish, but not in Spanish, poultry. In Bulgaria, quinolone AMR was also frequently observed, although their reported AMU did not follow the same trend.

Interestingly, we observed that country-wise estimated AMR richness significantly correlated between pig and poultry. Also, the countries with a high AMR richness were also the ones with higher AMR abundance (Italy, Spain, Bulgaria and Poland). While having higher abundance of the ubiquitous core genes, these countries had a higher number of unique genes in both pig and poultry production. The fact that countries’ AMR abundance and gene richness in pig and poultry tend to follow each other, could perhaps be explained with policy: if a country has strict AMU regulations in one livestock species, chances are that similar regulations are in place for other livestock species. Better host-separated, preferably herd-level, AMU data is needed to further explore this.

It has previously been reported that soil community composition structures the soil resistome.^26^ Using a wide array of environmental matrices, including fecal and waste water samples, this has also been shown for human habitats.^40^ We found the same to be true for pig and poultry resistomes, and additionally, we showed that the taxa-AMR association strength differs between countries. Horizontal gene transfer (HGT) could explain this phenomenon, if a larger proportion of certain countries’ resistome is mobile and AMR genes are more frequently introduced and re-introduced to genera. On the other hand, vertical AMR transmission can also play a role; if e.g. one country’s livestock is more isolated from trade. This would mean more livestock generations for the microbiome and resistome to diverge from those in other countries.

The determination of a set of core AMR genes for pig and poultry likely represents an underestimate of the true core resistomes. Requiring a certain percentage of samples to show evidence for a gene does not account for differential sequencing depth. The core genes discovered here thus likely represent a conservative subset of the true core resistomes.

In contrast to ResFinder, when using FRD we found no relationship between total drug use and total functional AMR abundance. This suggests that while many genes can provide AMR when cloned into e.g. *E. coli* in functional metagenomic assays, they might not provide AMR functionality in their natural hosts with natural expression levels. If most of them did, we could expect to see AM-based selection and an association to drug usage, like it is observed for the AMR genes in ResFinder - a database of genes known to provide AMR to their natural hosts. This finding echoes previous sentiments that one should carefully consider the risk to human health imposed by individual AMR genes.^41^ Some FRD genes might represent high risk, but we currently do not know what subset that is. Creating the FRD is a first step in trying to catalog the many AMR genes found in functional metagenomic studies. Screening sequenced pathogenic isolates and metagenomic assemblies for FRD genes, would be a good start for assessing their host range and risk potential.

The AMU data used in this study is not optimal. There is variation in drug use within each country’s farms that we did not account for by using a national average and we are unable to accurately distinguish between drugs used in different livestock species. Moreover the PCU denominator used by ESVAC may vary greatly between countries with differences in livestock populations and imports/exports that contribute to the biomass as calculated. Furthermore, the integrated herds enrolled in this study might represent only a limited subset of the overall livestock production in some countries. However, even with our crude AMU estimates, we found significant associations with total AMR abundance. The similar conclusion when considering the specific drug usage profile of each country indicates that the resistome is responding to AMU. The AMR-AMU association is well-documented for specific cultured indicator species and certain AM drugs, but relatively unknown when considering the whole microbiota and resistome and the newer approach of metagenomic shotgun sequencing.^3^,^8^ We do not know why the pig samples had a large within-country spread of total AMR, but perhaps the more heterogeneous production system and production management is responsible.

DNA extractions from the pooled poultry samples resulted in relatively low DNA yields. The protocol used was optimized for pig feces, human feces and sewage, but not poultry feces.^18^ The lower yields necessitated the use of a PCR-based library preparation kit, that can influence downstream analysis of shotgun sequencing.^42^. While the large difference between pig and poultry resistomes in our study is likely real, we caution the use of sensitive, quantitative analyses when comparing between samples prepared using different library preparation kits. For this reason, we have mostly tested within each reservoir. For future studies, one might consider using larger volumes of bird feces or otherwise optimize the protocol to ensure PCR-free library preparation is possible for all samples.

The sensitivity of metagenomic approaches does not yet rival phenotypic alternatives such as selective enrichment. We relied on read-mapping as opposed to a metagenomic assembly-based strategy for greater sensitivity. Still, there are AMR genes in important pathogens that we know are likely present but are below our detection limit. For example, we only found evidence for *bla*_CTX-M_ in three pig herds, whereas in phenotypic studies, the prevalence is high even among farms with no cephalosporin usage.^43^ Another tradeoff with the use of reference-based read mapping is our inability to identify mutated housekeeping genes granting AMR or distinguish between close homologs of the same AMR gene, e.g., the non-ESBL *bla*_TEM_ genes from those encoding ESBL variants.

The primary concern with read-mapping techniques, the lack of genomic context, can be solved using metagenomic assembly and binning approaches.^16^,^44^,^45^ In this way, AMR alleles in full length, their genomic context and their associated taxa have been identified in both pig, poultry and human fecal samples.^46^ As shown previously, the association between AMR and AMU is similar for metagenomics and traditional phenotypic methods, but several aspects make metagenomics an intriguing monitoring tool.^17^ The fact that both types of analyses (quantitative, sensitive read mapping and qualitative, context-giving binning) use the same raw data, makes metagenomics an attractive tool. In addition, the digital nature of shotgun data would also allow future re-use and form the basis of an invaluable historical archive, potentially usable for both AMR and pathogen tracking worldwide. While it is possible to identify thousands of AMR genes from environmental samples using functional metagenomics, and then track them using shotgun sequencing, their association to drug use and potential risk to human and animal health requires much further work to inform effective drug policies.

We found that the metagenomic resistome varied significantly in livestock, with large differences between the pig and poultry reservoir, but also within each livestock species, in a country-dependent manner. Within each country, we found different levels of variation, with some countries having more homogenous herds than others. Differences were seen both in total AMR abundance, but also abundances of AMR types and specific genes, including clinically relevant AMR genes. Some of this variation, we attributed to differential drug usage between the countries. We also identified the microbiome background as an important factor in determining the resistome in livestock, but found the strength of the association was country-dependent, at least in pigs. Interestingly, we found that AMR richness in one livestock species in a country is linked to the abundance in another livestock species. Finally, we observed some indications that newly described AMR genes from functionally metagenomic studies, might not provide AMR functionality when expressed in their natural host, even though they have the potential at the right expression levels in the right organism.

## Data availability

The DNA sequences (reads) from the 363 metagenomic samples from the 359 herds are deposited in the European Nucleotide Archive (ENA) under the project accession number PRJEB22062.

## Acknowledgements

We would like to thank all the anonymous pig and poultry herd owners who agreed to participate in the study and especially everyone involved in sampling and lab work: BE: Marjolijn Schlepers. BG: Teodora Ivanova, Nikolay Cholakov, Eva Gurova-Mehmedova, Krasen Penchev. DE: Franziska Nienhaus. DK: Cecilie Liv Nielsen, Pia Ryt-Hansen, Bonnie Hølstad, Bettina Rasmussen, Kirstine Nielsen. FR: Coton Jenna, Dorenlor Virginie, Eono Florent, Eveno Eric, Le Bouquin Sophie, Leon Denis, Thomas Rodolphe. IT: Mario Gherpelli, Marco Pegoraro, Virginia Carfora. NL: Daisy de Vries. PL: Beata Gawlik, Dorota Krasucka, Andrzej Hoszowski. The EFFORT project (www.effort-against-amr.eu) and the work presented here is supported by EU, FP7-KBBE-2013-7, grant agreement 613754.

## Author contributions

FMA, DH, JAW, TH, DM and the EFFORT group designed the study. FMA and BEK detailed the sampling and sequencing. ASRD and the EFFORT group carried out sampling. BEK and SJP conducted the DNA purification and organized with PM the sequencing. RBH, OL and TNP created the read-mapping pipeline. ER created the functional resistome database. PM, OL, RECL, LC, LAMS, HS, AB and HV carried out the bioinformatics and statistical analysis. PM created the figures and drafted the manuscript. All authors helped review, edit and complete the manuscript.

## Competing interests

The authors declare no competing financial interests.

## References

1. Who. ANTIMICROBIAL RESISTANCE Global Report on Surveillance 2014. World Heal. Organ. (2014).

2. Aarestrup, F. M. The livestock reservoir for antimicrobial resistance: a personal view on changing patterns of risks, effects of interventions and the way forward. Philos. Trans. R. Soc. Lond. B. Biol. Sci. 370, 20140085 (2015).

3. Chantziaras, I., Boyen, F., Callens, B. & Dewulf, J. Correlation between veterinary antimicrobial use and antimicrobial resistance in food-producing animals: A report on seven countries. J. Antimicrob. Chemother. 69, 827–834 (2014).

4. Dunlop, R. H. et al. Associations among antimicrobial drug treatments and antimicrobial resistance of fecal Escherichia coli of swine on 34 farrow-to-finish farms in Ontario, Canada. Prev. Vet. Med. 34, 283–305 (1998).

5. Aarestrup, F. M., Seyfarth, A. M., Emborg, H., Pedersen, K. & Bager, F. Effect of Abolishment of the Use of Antimicrobial Agents for Growth Promotion on Occurrence of Antimicrobial Resistance in Fecal Enterococci from Food Animals in Denmark. Antimicrob. Agents Chemother 45, 2054–2059 (2001).

6. Agersø, Y. & Aarestrup, F. M. Voluntary ban on cephalosporin use in Danish pig production has effectively reduced extended-spectrum cephalosporinase-producing Escherichia coli in slaughter pigs. J. Antimicrob. Chemother. 68, 569–72 (2013).

7. Dutil, L. et al. Ceftiofur resistance in Salmonella enterica serovar Heidelberg from chicken meat and humans, Canada. Emerg. Infect. Dis. 16, 48–54 (2010).

8. Dorado-García, A. et al. Quantitative assessment of antimicrobial resistance in livestock during the course of a nationwide antimicrobial use reduction in the Netherlands. J. Antimicrob. Chemother. dkw308 (2016). doi:10.1093/jac/dkw308

9. Aarestrup, F. M. et al. Resistance to antimicrobial agents used for animal therapy in pathogenic-, zoonotic- and indicator bacteria isolated from different food animals in Denmark: a baseline study for the Danish Integrated Antimicrobial Resistance Monitoring Programme (DANMAP). APMIS 106, 745–770 (1998).

10. Bronzwaer, S. Harmonised monitoring of antimicrobial resistance in Salmonella and Campylobacter isolates from food animals in the European Union. Clin. Microbiol. Infect. 14, 522–533 (2008).

11. McEwan, S. A., Aarestrup, F. M. & Jordan, D. in Antimicrobial resistance in bacteria of animal origin 397–413 (ASM Press, 2006).

12. Veterinary Medicines Division. European Medicines Agency, European Surveillance of Veterinary Antimicrobial Consumption, 2014. (2016).

13. ECDC. EFSA. ENA. ECDC/EFSA/EMA first joint report on the integrated analysis of the consumption of antimicrobial agents and occurrence of antimicrobial resistance in bacteria from humans and food-producing animals. EFSA J. 13, 4006 (2015).

14. EFSA. Technical specifications on the harmonised monitoring and reporting of antimicrobial resistance in Salmonella, Campylobacter and indicator Escherichia coli and Enterococcus spp. bacteria transmitted through food 1. EFSA J. 10, 64 (2012).

15. Nordahl Petersen, T. et al. Meta-genomic analysis of toilet waste from long distance flights; a step towards global surveillance of infectious diseases and antimicrobial resistance. Sci. Rep. 5, 11444 (2015).

16. Xiao, L. et al. A reference gene catalogue of the pig gut microbiome. Nat. Microbiol. 1, 16161 (2016).

17. Munk, P. et al. A sampling and metagenomic sequencing-based methodology for monitoring antimicrobial resistance in swine herds. J. Antimicrob. Chemother. dkw415 (2016). doi:10.1093/jac/dkw415

18. Knudsen, B. E. et al. Impact of Sample Type and DNA Isolation Procedure on Genomic Inference of Microbiome Composition. mSystems 1, 64394 (2016).

19. Li, H. & Durbin, R. Fast and accurate short read alignment with Burrows-Wheeler transform. Bioinformatics 25, 1754–1760 (2009).

20. Zankari, E. et al. Identification of acquired antimicrobial resistance genes. J. Antimicrob. Chemother. 67, 2640–2644 (2012).

21. Trapnell, C. et al. Differential gene and transcript expression analysis of RNA-seq experiments with TopHat and Cufflinks. Nat. Protoc. 7, 562–78 (2012).

22. Fu, L., Niu, B., Zhu, Z., Wu, S. & Li, W. CD-HIT: Accelerated for clustering the next-generation sequencing data. Bioinformatics 28, 3150–3152 (2012).

23. Pehrsson, E. C., Forsberg, K. J., Gibson, M. K., Ahmadi, S. & Dantas, G. Novel resistance functions uncovered using functional metagenomic investigations of resistance reservoirs. Front. Microbiol. 4, 1–11 (2013).

24. Moore, A. M. et al. Pediatric fecal microbiota harbor diverse and novel antibiotic resistance genes. PLoS One 8, (2013).

25. Sommer, M. O. a, Dantas, G. & Church, G. M. Functional characterization of the antibiotic resistance reservoir in the human microflora. Science 325, 1128–31 (2009).

26. Forsberg, K. J. et al. Bacterial phylogeny structures soil resistomes across habitats. Nature 509, 612– 6 (2014).

27. Legendre, P. & Gallagher, E. D. Ecologically meaningful transformations for ordination of species data. Oecologia 129, 271–280 (2001).

28. Veterinary Medicines Division. European Medicines Agency, European Surveillance of Veterinary Antimicrobial Consumption, 2016. (2016).

29. Jari Oksanen, F. Guillaume Blanchet, Michael Friendly, Roeland Kindt, Pierre Legendre, D., McGlinn, Peter R. Minchin, R. B. O'Hara, Gavin L. Simpson, Peter Solymos, M. H. H. & Stevens. vegan: Community Ecology Package. (2016).

30. Pielou, E. C. The measurement of diversity in different types of biological collections. J. Theor. Biol. 13, 131–144 (1966).

31. Paradis, E., Claude, J. & Strimmer, K. APE: Analyses of phylogenetics and evolution in R language. Bioinformatics 20, 289–290 (2004).

32. Letunic, I. & Bork, P. Interactive Tree Of Life (iTOL): An online tool for phylogenetic tree display and annotation. Bioinformatics 23, 127–128 (2007).

33. Wickham, H. ggplot2: Elegant Graphics for Data Analysis. (Springer-Verlag New York, 2009).

34. Bates, D., Mächler, M., Bolker, B. & Walker, S. Fitting Linear Mixed-Effects Models using lme4. eprint arXiv:1406.5823 67, 51 (2014).

35. Love, M. I., Huber, W. & Anders, S. Moderated estimation of fold change and dispersion for RNA-seq data with DESeq2. Genome Biol. 15, 550 (2014).

36. McMurdie, P. J. & Holmes, S. Waste not, want not: why rarefying microbiome data is inadmissible. PLoS Comput. Biol. 10, e1003531 (2014).

37. Hochberg, Y. & Benjamini, Y. Controlling the False Discovery Rate?: a Practical and Powerful Approach to Multiple Testing. 57, 289–300 (1995).

38. H Hasman, AM Hammerum, F Hansen, RS Hendriksen, B Olesen, Y Agersø, E Zankari, P Leekitcharoenphon, M Stegger, RS Kaas, LM Cavaco, DS Hansen, FM Aarestrup, R. S. Detection of Mcr-1 Encoding Plasmid-Mediated Colistin-Resistant Escherichia Coli Isolates From Human Bloodstream Infection and Imported Chicken Meat, Denmark 2015. Euro surveill 20, pii=30085 (2015).

39. Wang, Y. et al. A novel gene, optrA, that confers transferable resistance to oxazolidinones and phenicols and its presence in Enterococcus faecalis and Enterococcus faecium of human and animal origin. J. Antimicrob. Chemother. 70, 2182–2190 (2015).

40. Sun, M. et al. Interconnected microbiomes and resistomes in low-income human habitats SI. Nat. SI 61, 5985–5991 (2016).

41. Martínez, J. L., Coque, T. M. & Baquero, F. What is a resistance gene ? Ranking risk in resistomes. Nat. Publ. Gr. 13, 116–123 (2014).

42. Bowers, R. M. et al. Impact of library preparation protocols and template quantity on the metagenomic reconstruction of a mock microbial community. BMC Genomics 16, 856 (2015).

43. Hammerum, A. M. et al. Characterization of extended-spectrum β-lactamase (ESBL)-producing Escherichia coli obtained from Danish pigs, pig farmers and their families from farms with high or no consumption of third-or fourth-generation cephalosporins. J. Antimicrob. Chemother. 1–8 (2014). doi:10.1093/jac/dku180

44. Nielsen, H. B. et al. Identification and assembly of genomes and genetic elements in complex metagenomic samples without using reference genomes. Nat. Biotechnol. 32, (2014).

45. Anantharaman, K. et al. Thousands of microbial genomes shed light on interconnected biogeochemical processes in an aquifer system. Nat. Commun. 7, 1–11 (2016).

46. Ma, L. et al. Metagenomic Assembly Reveals Hosts of Antibiotic Resistance Genes and the Shared Resistome in Pig, Chicken, and Human Feces. Environ. Sci. Technol. 50, 420–427 (2016).

